# Fluorescence Properties of Collagen Types I-V: A Comprehensive Study of Spectral and Lifetime Characteristics

**DOI:** 10.64898/2026.02.21.706506

**Authors:** Xuhui Liu, Alexandra C. Adams, Xiangnan Zhou, Julien Bec, Laura Marcu

## Abstract

**Significance:** Collagen autofluorescence provides valuable intrinsic contrast for assessing tissue structure, composition, and pathology. However, a comprehensive understanding of the fluorescence properties across different collagen types remains limited. This knowledge gap may limit the development of advanced label-free fluorescence spectroscopy and imaging techniques for specific tissue characterization and diagnostic applications.

**Aim:** This study aims to comprehensively characterize the fluorescence intensity excitation-emission matrices (I-EEMs) and time-resolved excitation-emission matrices (TR-EEMs) of collagen standards from Types I, II, III, IV, and V obtained from various organ sources under both dry and hydrated conditions, to identify optimal excitation-emission parameters for each collagen type discrimination, and to establish a reference dataset that supports future research in label-free tissue characterization.

**Approach:** We employed a time-resolved fluorescence spectroscopy system equipped with an optical parametric oscillator laser (excitation: 200-2000 nm, pulse width: 30 ps) as an excitation source to generate I-EEMs and TR-EEMs of human and bovine collagen Types I-V. The fluorescence light was obtained by a multichannel plate photomultiplier tube through a monochromator (spectral range: 200-1000 nm). Measurements were conducted using collagen standards, under both dry and hydrated states. Additionally, photobleaching effects were assessed to ensure the reliability and reproducibility of fluorescence data.

**Results:** Each collagen type exhibited distinct I-EEM and TR-EEM signatures, with fluorescence lifetimes ranging from 2.5 ns (Type III, bovine skin) to 5.3 ns (Types II and V). Fibrillar collagens (Types I and V) displayed broader I-EEMs, whereas basement membrane collagen (Type IV) showed the narrowest spectral distribution. Organ-source-dependent variations were evident within the same collagen type. Type I collagen from human placenta exhibited an inverse lifetime–emission wavelength relationship compared to bovine sources. Hydration consistently red-shifted emission peaks into the 395–420 nm range and reduced fluorescence lifetimes across all collagen types (e.g., Type I bovine Achilles tendon: 3.2–5.0 ns dry vs. 3.0–4.5 ns hydrated). Despite excitation wavelength- and fluence-dependent photobleaching of fluorescence intensity, fluorescence lifetimes remained relatively stable, confirming the robustness of lifetime-based measurements.

**Conclusions:** This study establishes a comprehensive reference dataset for the fluorescence properties of collagen Types I-V and demonstrates the potential of combined I-EEMs and TR-EEMs analysis for tissue characterization. The results highlight species-, organ-, type-, and environment-specific optical fingerprints of similar collagens, which must be considered before implementing more in-depth studies on how the optical properties of collagen change in different medical applications.

## 1 Introduction

Collagen, the most abundant protein in mammals, is a key structural component of the extracellular matrix (ECM), providing mechanical strength and organization to tissues such as skin, bone, cartilage, tendons, and blood vessels [1]. Of the 28 known human collagen types, Types I through V are the most prevalent and functionally significant [2]. These five collagen types differ in amino acid composition, structural organization, tissue distribution, and biological function (Table 1).

**Table 1.** Properties of collagen Types I–V.

| Collagen Type | Structure | Primary Locations | Key Functions |
| --- | --- | --- | --- |
| Type I | Fibrillar | Skin, bone, tendons | Tensile strength and elasticity; >90% of total body collagen [3]. |
| Type II | Fibrillar | Cartilage | Cartilage integrity and joint flexibility[2] |
| Type III | Fibrillar | Skin, blood vessels, organs | Structural framework; synergistic with Type I in fibrillogenesis[4] |
| Type IV | Supramolecular network | Basement membranes | Forms the basement membrane scaffold. Filtration and cell adhesion[5] |
| Type V | Fibrillar | Cornea, placenta, hair | Regulation of collagen fibril formation and organization[6] |

Collagen’s characteristic triple-helical structure, rich in glycine, proline, and hydroxyproline, not only governs its biomechanical properties but also underlies its intrinsic optical properties [1]. Autofluorescence arises from various fluorescence species within collagen, including aromatic amino acids such as tyrosine and phenylalanine, as well as from natural covalent cross-links formed during tissue maturation and from advanced glycation end-products (AGEs) generated over time [7]. These endogenous fluorophores and cross-links, which are particularly abundant in long-lived collagens, impart unique spectral and lifetime signatures that can be exploited for non-invasive, label-free imaging.

Collagen autofluorescence has enabled differentiation between normal and pathological conditions in tissue and early detection of diseases such as fibrosis, atherosclerosis, and intervertebral disc degeneration [8–10]. Tyrosine and phenylalanine exhibit excitation maxima around 275 nm and 250 nm, respectively, with emission peaks at ∼300 nm and ∼280 nm [11]. In the visible range (390-450 nm), fluorescence is largely attributed to natural cross-links such as pyridinoline and deoxypyridinoline, as well as AGEs including pentosidine, which accumulate during aging or pathological states.

Fluorescence intensity excitation-emission matrices (I-EEMs) are shown to provide comprehensive spectral “fingerprints,” enabling identification of endogenous fluorophores by their unique excitation-emission profiles [12]. I-EEMs data has been used in the diagnosis of colonic cancer, cartilage defects, and in evaluation of extracellular matrix composition [13–15]. However, the potential contribution of time-resolved (TR) measurements of fluorescence decay or lifetime in conjunction with I-EEMs has not been systematically explored. TR-EEMs offers a more robust and multidimensional approach to tissue characterization, applicable to both non-pathologic and pathological contexts. Unlike fluorescence intensity, fluorescence lifetime is an intrinsically radiometric property that is largely independent of fluorophore concentration, illumination intensity, and excitation-collection geometry, making it a particularly reliable metric under heterogeneous or non-uniform illumination conditions [16].

Despite decades of investigations, collagen fluorescence studies largely have been restricted to single or limited of excitation wavelengths or to individual collagen types, without accounting for type-specific differences, hydration effects, or organ-source variability [17–19]. In particular, comparative lifetime analyses across collagen standards from Types I–V under diverse experimental conditions and environments, including the difference between the standard form and PBS hydrated form, remains scarce.

While preliminary evidence suggests that water molecules may quench collagen fluorescence [20], comprehensive time-resolved studies are lacking. Similarly, organ-specific comparisons (e.g., human skin vs. placenta; bovine skin vs. Achilles tendon) are rare, despite known differences in cross-linking density and mechanical properties that are expected to influence the fluorescence signatures [19].

This fragmented body of knowledge highlights an essential need: the lack of a comprehensive reference dataset characterizing the spectral and fluorescence lifetime properties of collagen Types I–V across varying hydration states and tissue origins. Establishing such a dataset would support the optimization of fluorescence-based diagnostic tools, inform the design of collagen-derived biomaterials, and advance understanding of tissue-specific collagen behavior in both healthy and diseased states.

To address this gap, we constructed a time-resolved fluorescence spectroscopy (TRFS) apparatus equipped with an optical parametric oscillator laser as a light source that enabled extensive characterization of collagen samples. The goals of this study are to: (1) Characterize I-EEMs and TR-EEMs for collagen Types I–V standards, identifying optimal excitation-emission parameters and establishing a reference dataset; (2) Assess the effect of hydration (dry vs hydrated) on the I-EEMs and TR-EEMs, elucidating water’s role in modulating fluorophore microenvironments; (3) Compare collagens from distinct organ sources—human skin vs. placenta and bovine skin vs. Achilles tendon—to identify tissue-specific fluorescence signatures. Additionally, we investigated the photobleaching impact on collagen’s fluorescence emissions and lifetimes from different organ sources under different fluence rates. Through this systematic approach, we aim to provide a standardized reference dataset for collagen autofluorescence properties to support biophysical, biochemical, and biomedical engineering.

## 2 Methods

### 2.1 Experimental apparatus

To characterize the emission spectra and lifetime of collagens, we developed an experimental setup consisting of a customized pulse-sampling fluorescence lifetime (psFLT) spectroscopic system, as illustrated in Fig. 1. The system employed an optical parametric oscillator (OPO) laser (tunable wavelength range: 200-2000 nm, pulse width: 30 ps, repetition rate: 1 kHz, PT403, EKSPLA) as an excitation light source. A power control module maintained constant pulse energy at 0.4 µJ throughout all measurements. This module comprised a rotatable half-wave plate (AHWP05M-580, Thorlabs), a polarizer (GT5A, Thorlabs), a beamsplitter (70/30 reflection/transmission ratio, UV-NIR, Edmund Optics), and a power meter (Ophir Optronics Solution). The excitation beam was focused using a spherical lens (f = 40 mm, AR coating: 350-700 nm, LBF254-040, Thorlabs) and coupled into a bifurcated fiber probe (Polymicro Technologies LLC, Phoenix, Arizona). The probe was custom designed. It consisted of a central illumination fiber (600 μm) that was tapered to a distal core diameter of 1025 μm. Thus, the NA of the illumination channel was ∼0.11. The central fiber was surrounded concentrically by a ring of 18 collection fibers, each of 200-μm core size. The SDS, which accounts for the cladding and jacket sizes and the intervening epoxy, was 0.64 mm. Both the illumination and the collection paths of the probe were 2 m long.

**Fig. 1.**
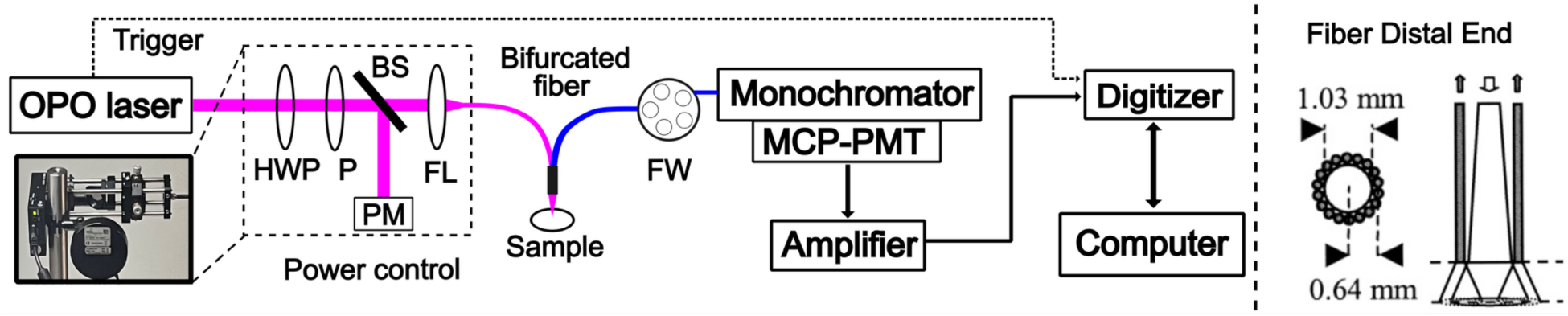
Instrumental apparatus schematic and the cross-sectional and side view of the bifurcated fiber. HWP: half-wave plate, P: polarizer, PM: power meter, BS: Beamsplitter, FL: focal lens, FW: filter wheel. MCP-PMT: multichannel plate photomultiplier tube.

Fluorescence emission from the sample was collected through the same bifurcated fiber probe and spectrally resolved using a monochromator (spectral resolution: 0.3 nm, MicroHR, Horiba). To eliminate back-reflected excitation light, a filter wheel containing five long-pass filters (cut-on wavelengths: 325, 355, 370, 395, and 450 nm, Edmund Optics) was positioned in the emission path.

Fluorescence signals were detected using a multichannel plate photomultiplier tube (MCP-PMT, spectral response: 160-850 nm, FWHM: 45 ps, R3809U-50, Hamamatsu) powered by a 0-2400 V supply (Ortec 556). The MCP-PMT output was amplified by a high-speed amplifier (gain: 40 dB; bandwidth: 0.01–3000 MHz; AM-1607-SMA, Miteq) prior to digitization. Time-resolved fluorescence emission was recorded using a high-speed digitizer (1.5 GHz, 5 GS/s, PIXe-5162, National Instruments) synchronized to TTL trigger pulses from the OPO laser.

### 2.2 OPO-psFLT system performance evaluation: Experimental protocol

The performance of the custom OPO-psFLT system in accurately resolving fluorescence decays was validated using reference dyes with well-characterized emission spectra and lifetimes. Solutions of POPOP and Coumarin 460 (Sigma-Aldrich) were prepared at 100 µM concentration in ethanol. EEMs were acquired over 300-400 excitation and 350-500 nm emission ranges for POPOP and 330-450 excitation and 400-520 nm emission ranges for Coumarin 460, with 10 nm wavelength step. A minimum spectral separation of 15 nm between excitation and emission wavelengths was enforced via a custom monochromator control software to prevent excitation light leakage. The excitation pulse energy was maintained at 0.4 µJ across all wavelengths using a power control module (Fig. 1).

### 2.3 Collagen Sample Preparation

Five commercially available collagen types (I-V) from various species and organ sources were investigated, as summarized in Table 2. For dry-form measurements, the collagen powders were placed on glass coverslips. For hydrated measurements, dry collagen samples were thoroughly mixed with phosphate-buffered saline (pH 7.4, PBS) and allowed to equilibrate for 5 minutes prior to data acquisition. All measurements were conducted at room temperature.

**Table 2:** Collagen samples used in this study.

| Collagen Type | Organ Source | Measurement Form |  |
| --- | --- | --- | --- |
| Type I | Human skin | Dry | Hydrated |
|  | Human placenta | Dry | Hydrated |
|  | Bovine Achilles tendon | Dry | Hydrated |
|  | Bovine skin | Dry | --- |
| Type II | Bovine nasal septum | Dry | Hydrated |
| Type III | Human placenta | Dry | --- |
| Type IV | Human placenta | Dry | --- |
| Type V | Bovine Achilles tendon | Dry | Hydrated |

### 2.4 Experimental Procedures

#### 2.4.1 Time-resolved excitation-emission matrix (TR-EEM) measurements

TR-EEM characterizations were performed on all collagen samples listed in Table 2. All samples were initially measured in dry state and selected collagen types (I, II, and V) were additionally evaluated in hydrated state. Based on published collagen absorption and emission spectra [7], laser excitation wavelengths spanning 315-405 nm (10 nm increment) were used, while fluorescence were collected over 350-500 nm (5 nm increment). The laser irradiation time per emission wavelength collection was 2 seconds, resulting in a total acquisition time of approximately 900 s for a complete EEM measurement, including laser power adjustments and wavelength transitions. Excitation pulse energy was maintained at 0.4 µJ throughout all measurements.

#### 2.4.2 Photobleaching Assessment

Given the extended laser exposure during EEM measurements, photobleaching effects were systematically evaluated on Type I collagen from bovine Achilles tendon and skin. Samples were excited at 355 nm and monitored continuously for 900 seconds at three emission wavelengths: 380, 405, and 450 nm for Achilles tendon, and 390, 425, and 460 nm for bovine skin. These wavelengths were selected based on the characteristic Type I collagen emission spectra to assess photobleaching at the emission peak and adjacent spectral regions [7, 21].

Two excitation pulse energies (0.4 and 0.8 µJ), corresponding to fluence rates of 50 µW/mm² and 100 µW/mm², were tested. The total fluence delivered over the measurement period was 45 mJ/mm^2^ and 90 mJ/mm^2^, respectively. Laser pulse-to-pulse energy variability was less than 5%. Five replicate measurements were performed for each condition, with each measurement taken on a fresh sample location.

Additional photobleaching experiments were conducted to assess wavelength-dependent effects using three excitation wavelengths (335, 355, and 375 nm) at constant pulse energy (0.4 µJ) on Type I collagen from bovine Achilles tendon. To evaluate the impact of photobleaching on emission intensity and fluorescence lifetime, repeated measurements were performed on the same sample site at each excitation wavelengths, with emission collected from 360-450 nm in 10 nm increments.

### 2.5 Data Analysis and Visualization

Fluorescence decay curves and average lifetimes were computed using a fast algorithm (<10 µs per decay) constrained least-squares deconvolution algorithm based on the Laguerre expansion method [22]. Fluorescence intensity was calculated by integrating the area under the curve (AUC) under the fluorescence decay curve.

To visualize the fluorescence intensity and lifetime characteristics at different excitation and emission wavelengths, we generated I-EEMs maps, TR-EEMs maps, emission spectra, and lifetime spectra for each measured sample. The I-EEMs map was generated by normalizing the individual fluorescence intensity to the peak intensity across all excitation-emission combinations, while the TR-EEMs map was generated by calculating the average lifetimes from each excitation-emission wavelength and grouping them as an excitation-emission map. For each excitation wavelength, the emission-spectra were generated by normalizing to its peak emission, while the lifetime-spectra were generated by calculating the average lifetime at each emission wavelength. Intensity and lifetime spectra for multiple excitations were plotted together to show the excitation-dependent characteristics. The fluorescence intensities that were lower than the minimal threshold for deconvolution were excluded from analysis [23].

## 3 Results

### 3.1 Evaluation of the OPO-psFLT System Performance on Fluorescence Standard Dyes

The I-EEMs and TR-EEMs maps of the standard fluorescence dyes are depicted in Fig. 2a and 2d. POPOP exhibited a peak emission at 420 nm when excited at 350 nm, whereas Coumarin 460 (C460) demonstrated peak emission at 450 nm under 380 nm excitation. For both dyes, normalized intensity and lifetime spectra remained consistent exhibited complete overlap across different excitation wavelengths (Fig. 2b and 2e). The TR-EEMs and lifetime spectra revealed mean fluorescence lifetimes of approximately 1.5 ns for POPOP and 3.1 ns for C460 across most excitation–emission combinations, including those with low emission intensity (Fig. 2a and 2b). These measurements agree well with published values of 1.43 ns for POPOP and 3.2 ns for C460 [24, 25]. Representative waveforms are presented in Fig. 2c (POPOP, 400 nm emission, 350 nm excitation) and Fig. 2f (C460, 450 nm emission, 360 nm excitation). In both cases, normalized residuals remained within the confidence intervals and exhibited no autocorrelation structure, confirming the robustness of the deconvolution approach.

**Fig.2.**
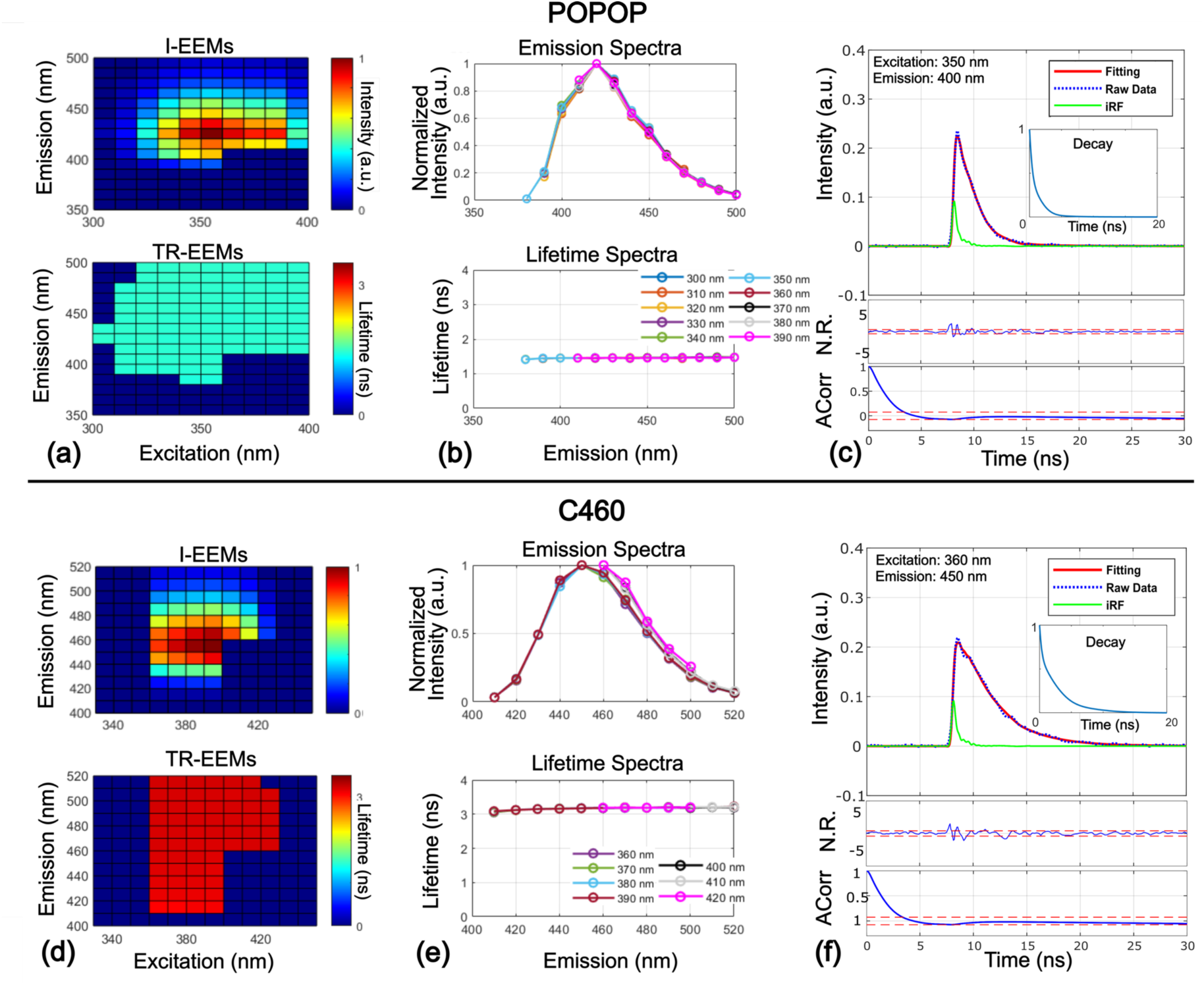
OPO-FLIm system validation using fluorescence standard dyes for POPOP (panel set: a, b, c) and C460 (panel set: d, e, f). I-EEMs and TR-EEMs maps are shown in (a) and (d). Emission-spectra (upper) and lifetime-spectra (lower) are shown in (b) and (e). Representative plots of raw data, fitting data, instrument response function (iRF), and deconvolved decay with normalized residuals (N. R.) and autocorrelation (Acorr.) are shown in (c) and (f).

### 3.2 Excitation-Emission Matrices and Fluorescence Lifetime Characterization

#### 3.2.1 Type I Collagen in dry form

Type I collagen from different organ sources exhibited distinct I-EEM and TR-EEM characteristics (Fig. 3a to 3d). Peak excitation-emission wavelengths were identified at 345/405 nm for bovine Achilles tendon, 325/405 nm for bovine skin, 315/400 nm for human skin, and 335/395 nm for human placenta. Emission spectra for all Type I collagen sources demonstrated red-shifted peaks with increasing excitation wavelengths. TR-EEMs maps and lifetime spectra revealed relatively constant lifetime distributions across excitation-emission combinations for bovine Achilles tendon, bovine skin, and human skin samples, even for lower fluorescence intensities. Both emission wavelength and tissue type exhibited distinct impacts on fluorescence lifetimes. Specifically, red-shifted emission wavelengths corresponded to longer lifetimes for collagen from bovine Achilles tendon, bovine skin, and human skin, whereas human placenta collagen exhibited decreased lifetimes with increasing emission wavelengths. Bovine Achilles tendon collagen exhibited the longest lifetimes (3.2–5.0 ns), while bovine skin collagen displayed the shortest lifetimes (2.9–4.0 ns).

**Fig.3.**
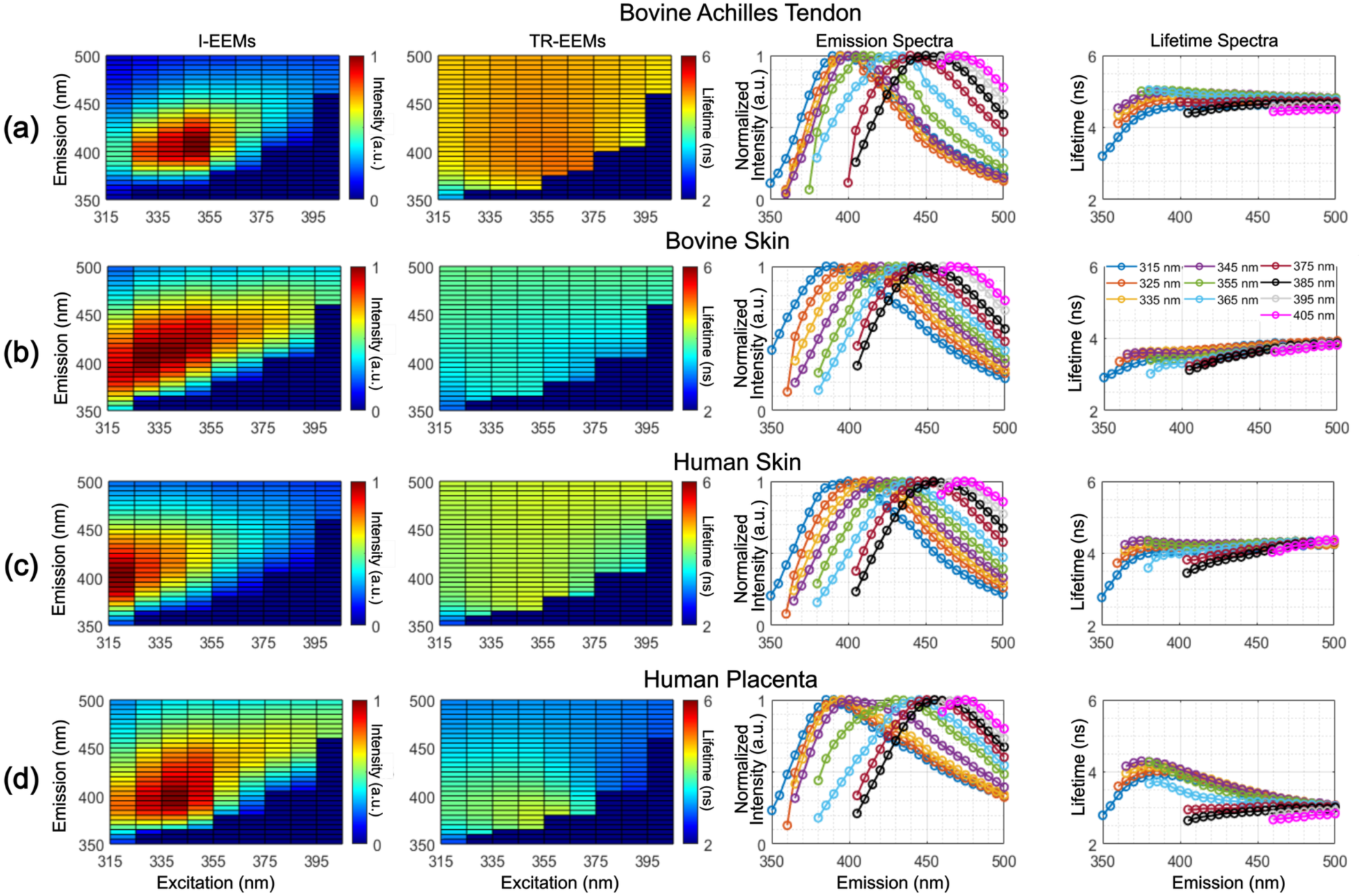
Fluorescence properties of Type I collagens in dry form. (a) to (d) From left to right: I-EEMs map, TR-EEMs map, emission-spectra for all excitation wavelengths, and lifetime-spectra across different excitations for Type I collagen from bovine Achilles tendon, bovine skin, human skin, and human placenta, respectively.

Hydration significantly altered the fluorescence properties of Type I collagens from bovine Achilles tendon, human skin, and human placenta (Fig. 4a to 4c). For example, the fluorescence peak of Type I collagen from bovine Achilles tendon centered at 345/405 nm in the dry form, shifted to 325/415 nm upon hydration, corresponding to a 20 nm blue shift in excitation and 10 nm red shift in emission, demonstrated a broader I-EEM. The FWHM of emission spectra also increased by hydration. For example, when the Type I collagen from bovine Achilles tendon was excited at 315 nm (blue curve), the FWHM of its emission spectra increased by 28% nm (70 nm to 90 nm) due to hydration. Similar trends were observed in collagens from other organ sources. Hydration decreased fluorescence lifetimes while maintaining similar patterns across excitation and emission wavelengths. TR-EEMs and lifetime spectra decreased consistently across all hydrated Type I collagen samples. For example, Achilles tendon collagen lifetimes ranged from 3.2–5.0 ns in dry form and 3.0–4.5 ns in hydrated form. The magnitude of lifetime changes remained similar across emission wavelengths and organ sources compared with dry collagen.

**Fig. 4.**
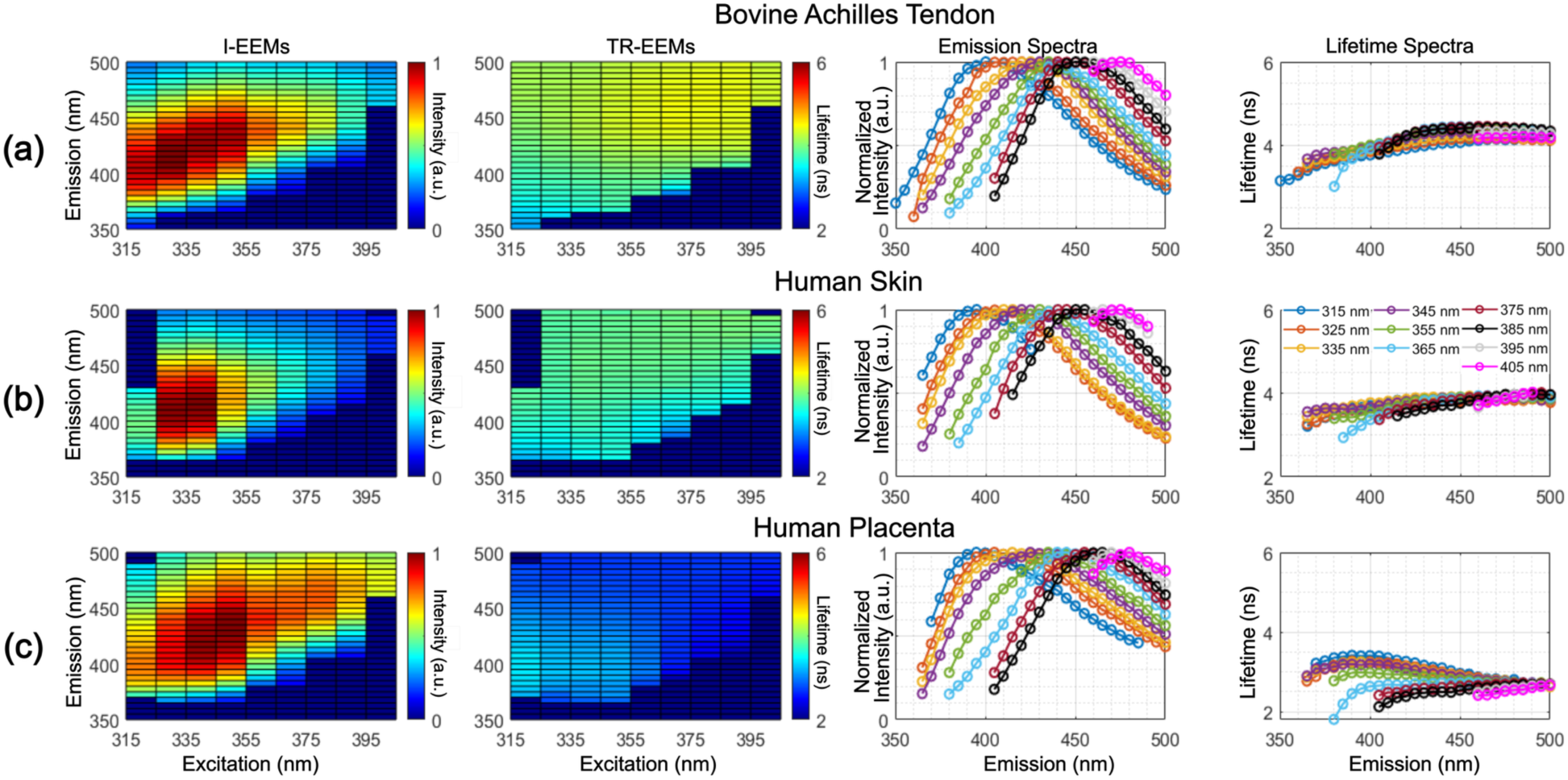
Fluorescence properties of hydrated Type I collagens. (a) to (c) From left to right: I-EEMs map, TR-EEMs map, emission-spectra, and lifetime-spectra across all excitation wavelengths for collagen sample from distinct sources: bovine Achilles tendon, human skin, and human placenta, respectively.

#### 3.2.2 Type II Collagen in dry and hydrated form

The fluorescence emission properties of Type II collagen from bovine nasal septum in both dry and hydrated forms are depicted in Figure 5a and Figure 5b, respectively. Hydrated samples exhibited I-EEMs with peak intensity at 325/395 nm excitation-emission wavelength, compared to 315/385 nm for dry samples, corresponding to a 10 nm red shift in excitation and emission. TR-EEMs and lifetime spectra ranged from 3.9 to 5.3 ns in dry Type II collagen and from 3.4 to 4.9 ns in hydrated samples. The shortest (3.9 ns) and longest (5.3 ns) lifetimes in dry collagen occurred at 315/350 nm and 355/395 nm, respectively. For hydrated collagen, the shortest (3.4 ns) and longest (4.9 ns) lifetimes were observed at 315/355 nm and 355/435 nm, respectively. The lifetime spectra exhibited distinct trends between dry and hydrated sample. In hydrated samples, the lifetime increased monotonically from 3.4 to 4.9 ns with increasing emission wavelength, whereas in dry sample, the lifetime initially increased then deceased with the increasing emission wavelength. Additionally, hydrated sample shows greater variations in lifetime spectra across excitation wavelength.

**Fig. 5.**
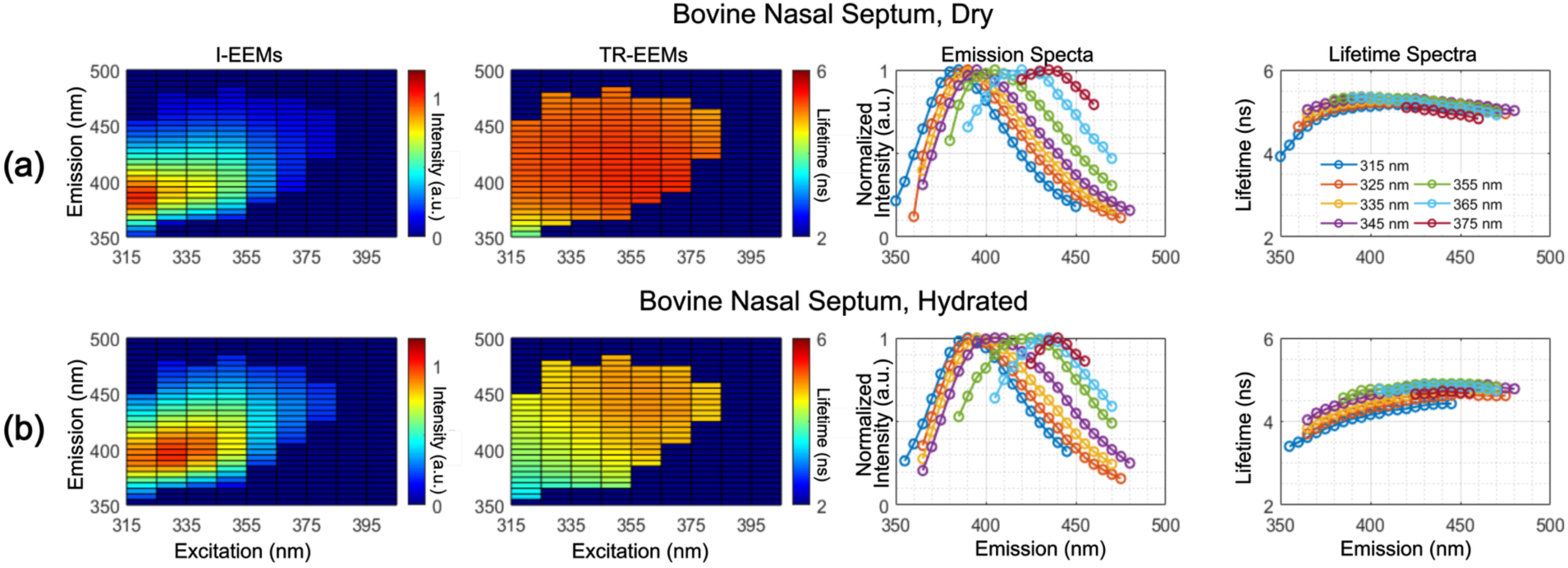
Fluorescence properties of Type II collagen from bovine nasal septum. (a) and (b) From left to right: I-EEMs map, TR-EEMs map, emission-spectra, and lifetime-spectra across all excitation wavelengths for dry and hydrated collagen, respectively.

#### 3.2.3 Type III Collagen in dry form

Type III collagens from bovine skin and human placenta were characterized in dry form, revealing tissue-type-dependent fluorescence signature (Fig. 6a and 6b). Bovine skin collagen exhibited broader I-EEMs with a peak intensity at 345/440 nm excitation-emission wavelength, whereas human placenta collagen peaked at 315/380 nm. The emission spectra also demonstrated tissue-type-dependent properties. For example, the peak emission of bovine skin collagen was at 440 nm when excited at 355 nm (green curve, Fig. 6a), while the peak emission shifted to 400 nm in human placenta collagen (green curve, Fig. 6b). TR-EEMs map and lifetime-spectra ranged from 2.5 to 4.3 ns for bovine skin collagen and from 3.0 to 4.9 ns for human placenta collagen, while depicts different trend as a function emission wavelength. The lifetime spectra showed different trends in tissue type. in bovine skin collagen, the lifetime increased monotonically with increasing emission wavelength, whereas in human placenta collagen, lifetime initially increased then deceased with the increasing emission wavelength. The shortest and longest lifetimes for bovine skin collagen (2.5 ns and 4.3 ns) occurred at 315/350 nm and 355/455 nm, respectively. For human placenta collagen, these extremes (3.0 ns and 4.9 ns) were found at 315/350 nm and 355/380 nm, respectively.

**Fig. 6.**
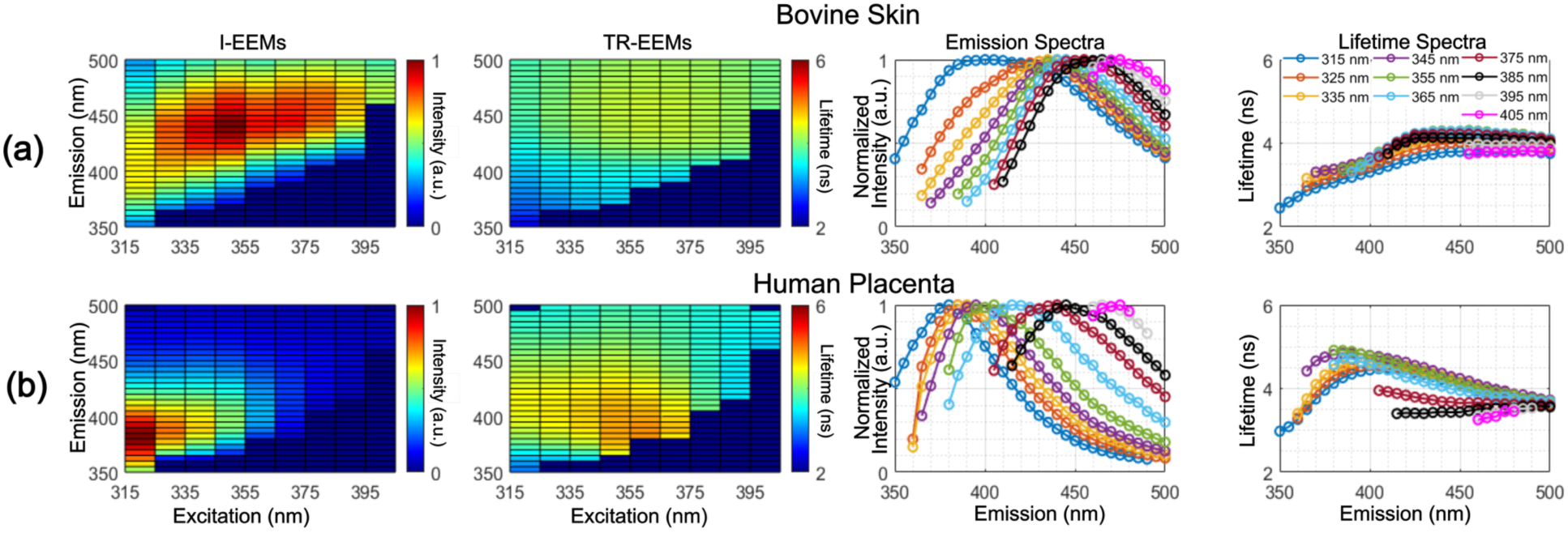
Fluorescence properties of Type III collagen in dry form. (a) and (b) From left to right: I-EEMs map, TR-EEMs map, emission-spectra, and lifetime-spectra across all excitation wavelengths for Type III collagen from bovine skin and human placenta, respectively.

#### 3.2.4 Type IV Collagen in dry form

The fluorescence emission properties of Type IV collagen from human placenta in dry form are depicted in Fig. 7. The I-EEMs revealed peak intensity at 315/380 nm excitation-emission wavelength. Emission spectra showed a red-shifted peak emission with the increase of excitation wavelength. TR-EEMs map and lifetime-spectra ranged from 4.1 to 5.1 ns across the measurement matrix, with higher values (∼5.1 ns) at 345/385 nm and lower values (∼4.1 ns) at 375/450 nm.

**Fig. 7.**
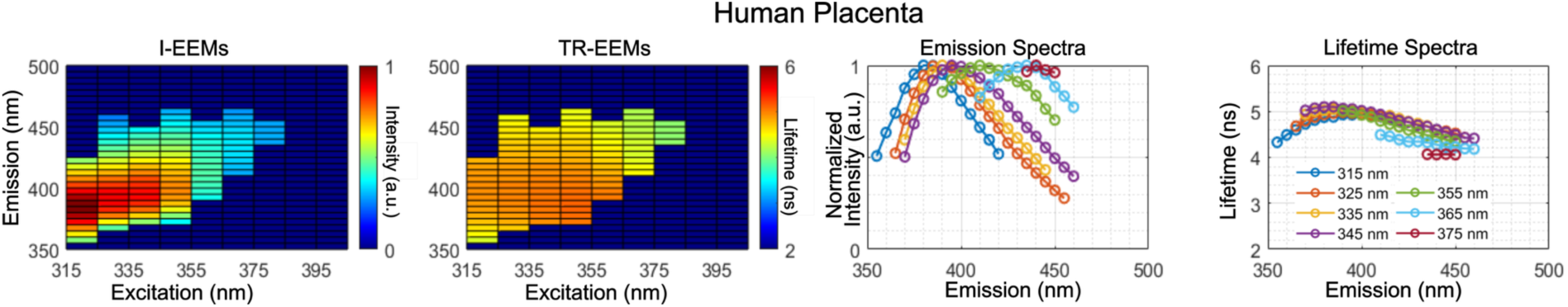
Fluorescence properties of Type IV collagen from human placenta in dry form. From left to right: I-EEMs, TR-EEMs map, emission-spectra, and lifetime-spectra across all excitation wavelengths.

#### 3.2.5 Type V Collagen

Type V collagen from bovine Achilles tendon was characterized in both dry and hydrated forms (Fig. 8a and 8b). Hydrated samples exhibited I-EEMs with peak intensity at 345/395 nm excitation-emission wavelength, compared to 335/400 nm for the dry samples, corresponding to a 10 nm red shift in excitation and a 5 nm blue shift in emission. TR-EEMs and lifetime spectra ranged from 3.3 to 5.3 ns in dry Type V collagen and from 3.3 to 4.9 ns in hydrated samples. The shortest (3.3 ns) and longest (5.3 ns) lifetimes in dry collagen occurred at 345/395 nm and 335/400 nm, respectively. For hydrated collagen, the shortest (3.3 ns) and longest (4.9 ns) lifetimes were observed at 315/355 nm and 355/435 nm, respectively. The lifetime spectra showed distinct trends between dry and hydrated sample. In dry samples, the lifetime initially increased from 3.3 to 5.3 ns and then reached at a plateau with increasing emission wavelength, whereas in hydrated samples, lifetime increased from 3.3 to 4.9 with increasing emission wavelength. Additionally, the hydrated sample shows greater variation in lifetime spectra across excitation.

**Fig. 8.**
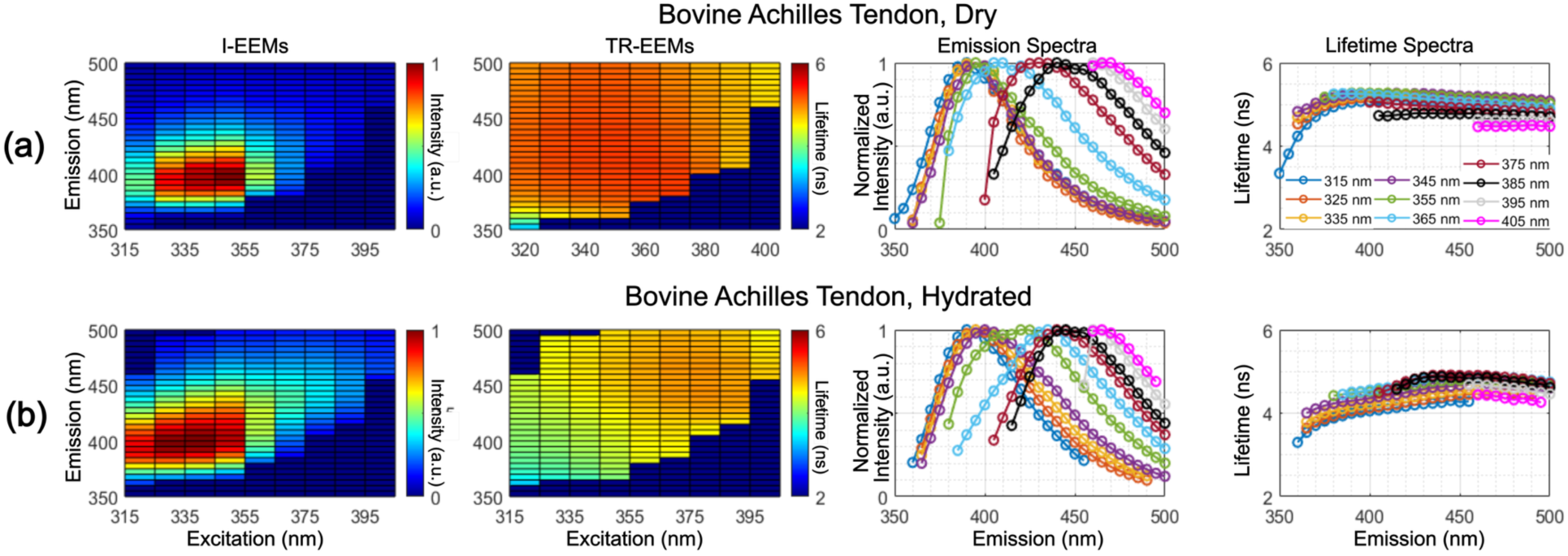
Fluorescence properties of Type V collagen from bovine Achilles tendon. (a) dry and (b) hydrated collagen samples; From left to right: I-EEMs, TR-EEMs, emission-spectra, and lifetime-spectra across all excitation wavelengths.

### 3.3 Photobleaching Effects on Fluorescence Properties

#### 3.3.1 Continuous Irradiation Studies

Photobleaching analysis of Type I collagen under continuous laser irradiation revealed wavelength and excitation energy-dependent emission intensity decay patterns (Fig. 9). For bovine Achilles tendon Type I collagen (Fig. 9a), fluorescence intensity decreased progressively with irradiation time. Half-decay time, when the intensity decreases to 50% of its maximum, was used to quantify photobleaching rate. At 380 nm emission, photobleaching effect was least pronounced, with half-decay times of 44 s and 24 s for pulse energies of 0.4 µJ and 0.8 µJ, respectively. Despite significant intensity decreases, fluorescence lifetimes values remained relatively constant throughout the irradiation period. Similar behavior was observed for bovine skin collagen (Fig. 9b), where emission at 425 nm exhibited the lowest sensitivity to photobleaching, with half-decay times of 56 s and 28 s for both energy levels. Fluorescence lifetimes again showed minimal variation with irradiation time. Excitation wavelength-dependent photobleaching was further assessed for bovine Achilles tendon collagen at 405 nm emission using 335, 355, and 375 nm excitation (Fig. 9c). Photobleaching was found most pronounced under 335 nm excitation and least under 355 nm excitation.

**Fig. 9.**
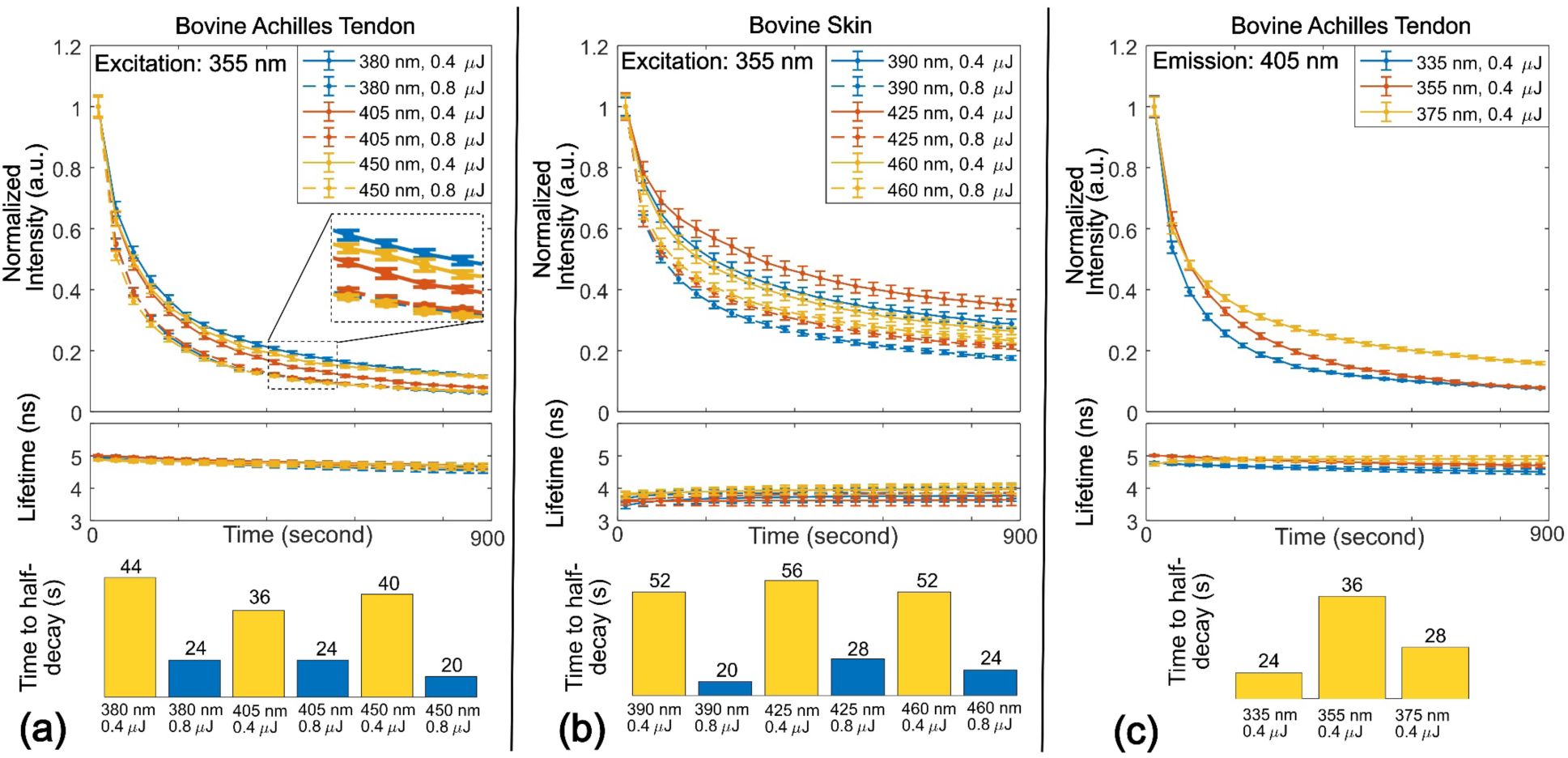
Collagen Type I photobleaching effect. (a) bovine Achilles tendon and (b) bovine skin: Behavior of the fluorescence intensity and lifetime for three emission wavelengths (i.e. 380 nm, 405 nm, and 450 nm) as a function of irradiation time for sole excitation wavelength (355 nm) and two pulse energy levels (0.4 µJ and 0.8 µJ). (c) Bovine Achilles tendon: Behavior of the fluorescence intensity and lifetime at for sole emission wavelength (i.e. 405 nm) as a function of irradiation time for three excitation wavelengths (i.e. 335, 355, and 375 nm).

#### 3.3.2 Impact of prolonged laser irradiation on fluorescence emission intensity-spectra and spectral shape

Sequential measurements on the same Type I collagen site (bovine Achilles tendon) revealed cumulative photobleaching effects over three measurement cycles (∼150 s per cycle). Significant intensity decrease was observed after repeated measurements (Fig. 10a-10c). Due to the fast-photobleaching rate at the beginning of irradiation, the normalized emission spectrum shows larger variations across 3 replicate measurements with 355 nm excitation. Fluorescence lifetimes values remained nearly constant across all measurement repeats.

**Fig. 10.**
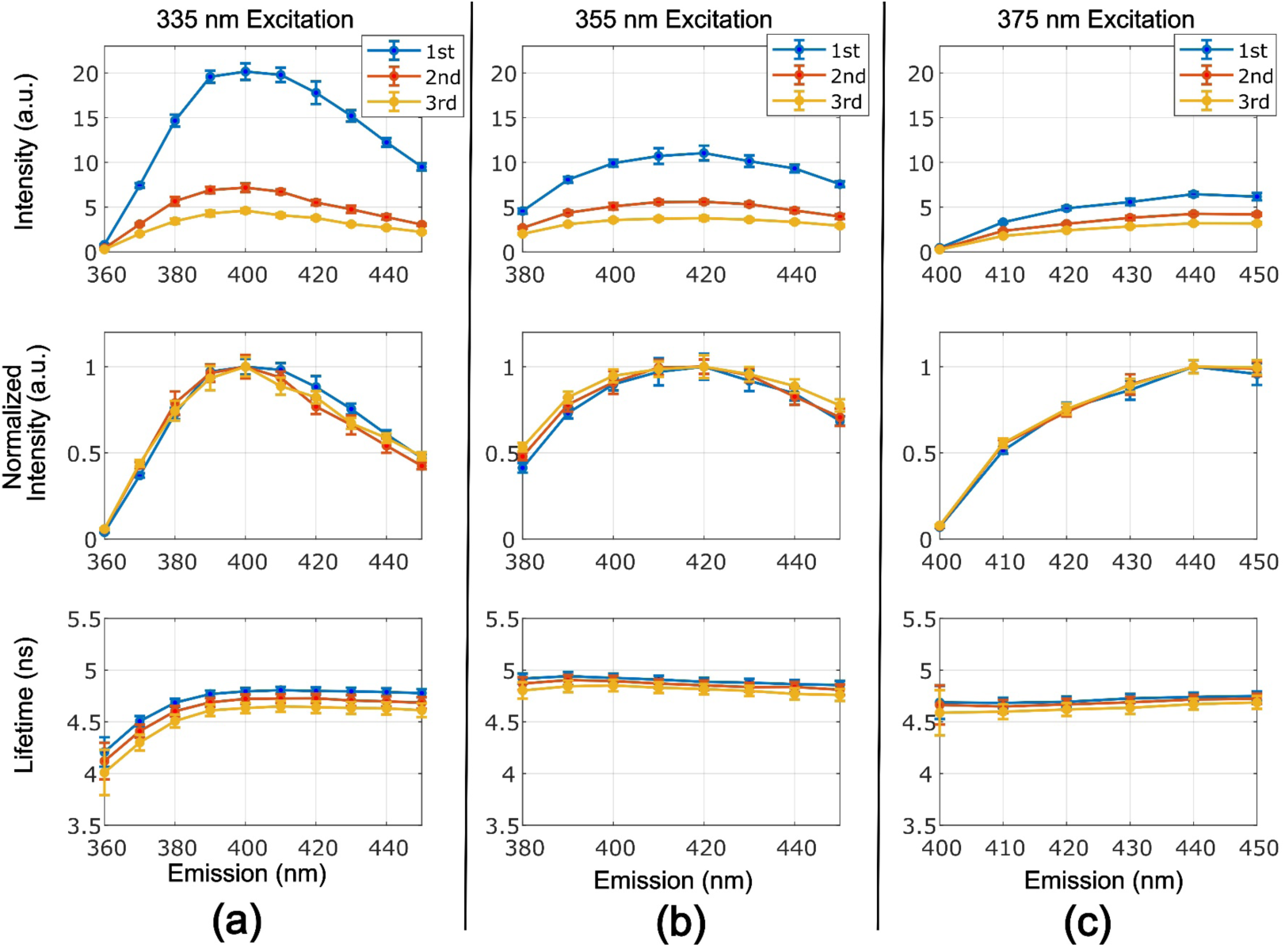
Variations of fluorescence emission (emission-spectra and spectral shape) and fluorescence lifetime for three repeated measurements. (a) to (c) From top to bottom: fluorescence intensity, normalized spectral shape, and lifetime spectra for excitation wavelengths at 335 nm, 355 nm, and 375 nm.

**Fig. 11.**
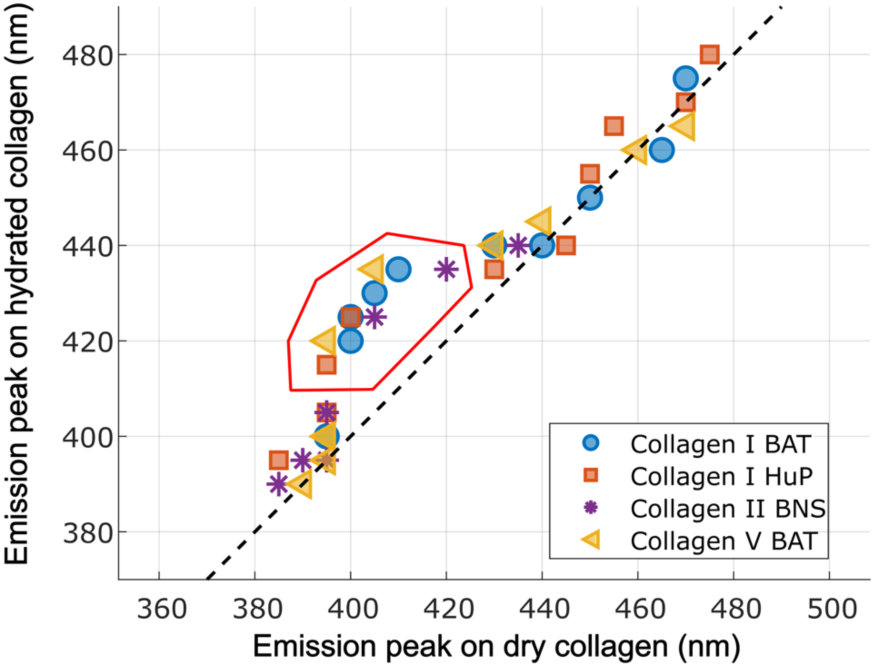
Impact of hydration and excitation wavelength on emission spectra. Scatter plot of emission peaks for dry versus hydrated collagen I, II, and V. Collagens with shifted emission peaks are highlighted with red marker. BAT: bovine Achilles tendon; HuP: human placenta; BNS: bovine nasal septum.

### 3.4 The Impact of Hydration on Fluorescence Emissions

Comparison of dry and hydrated states for collagen Types I, II, and V revealed consistent hydration-induced spectral shifts in specific excitation-emission spectral bands. Across all tested collagen types, hydration systematically red-shifted emission peaks to longer wavelengths within 395 - 420 nm range. Notably, this spectral shift pattern was observed for all collagens regardless of species or organ source.

## 4 Discussion

This study provides the first systematic characterization of both I-EEMs and TR-EEMs for collagen Types I–V samples across multiple organ sources and hydration states. The resulting dataset reveals that even in purified collagen samples, fluorescence emission signatures vary significantly, an important factor to consider when interpreting the tissue autofluorescence features in both research and clinical applications.

### 4.1 Tissue-Specific and Wavelength-Dependent Fluorescence Properties

Our data demonstrate that organ-source introduces substantial fluorescence variability across all collagen types. Type I collagen obtained from four different tissues exhibited peak excitation-emission wavelengths ranging from 315/400 nm (human skin) to 345/405 nm (bovine Achilles tendon), with corresponding lifetimes ranging from 2.9–4.0 ns (bovine skin) to 3.2–5.0 ns (bovine Achilles tendon). These organ-dependent differences indicate that tissue-specific factors such as cross-linking density and collagen maturation state [19], can modulate fluorescence properties to an extent comparable to collagen type itself. This highlights the importance of organ-specific reference for accurate tissue characterization. Across collagen types, the most prominent distinction was between fibrillar collagens (Types I and V), which displayed broad I-EEM profiles, and basement membrane collagen (Type IV), which exhibited the narrowest spectral distribution along with a lifetime range (4.1–5.1 ns). Moreover, excitation-wavelength-dependent shifts in emission spectra and fluorescence lifetimes underscore the photophysical complexity of collagen autofluorescence and support the presence of multiple emissive species, including aromatic amino acids and collagen cross-links [9, 26, 27].

### 4.2 Hydration-Induced Fluorescence Variations

Hydration generated two consistent effects across all collagen types and organ sources tested, specifically red-shifted emission peaks into the 395–420 nm range and decrease of fluorescence average lifetimes. For example, Type I collagen from bovine Achilles tendon lifetime has decreased from 3.2–5.0 ns (dry) to 3.0–4.5 ns (hydrated) while the emission FWHM has broaden by up to 28%. These changes are consistent with hydration-dependent modulation of excited-state relaxation pathways through water-protein hydrogen-bond networks [28], and enhanced non-radiative deactivation in aqueous environments [29]. The selective nature of these shifts, confined to specific excitation-emission bands, supports a multi-component model of collagen autofluorescence with fluorophores differing in solvent accessibility[30]. Since collagen is hydrated in vivo, dry-sample measurements should be interpreted as baseline references rather than surrogates for physiological conditions, and the paired characterization presented here provides reference standards for translating bench-top measurements to in vivo fluorescence lifetime imaging.

### 4.3 Photobleaching Effects and Their Implications for Fluorescence-Based Measurements

Photobleaching rates varied with collagen type and source, excitation wavelength, and pulse energy. Achilles tendon collagen photobleached more rapidly than skin collagen[31], and a clear excitation-dependent hierarchy emerged (335 nm > 375 nm > 355 nm), likely reflecting stronger overlap between 335 nm excitation and collagen cross-link absorption bands[32].

The central finding, however, is that fluorescence lifetimes remained relatively stable throughout prolonged irradiation despite substantial intensity reductions[21]. Because lifetime reflects intrinsic excited-state dynamics rather than fluorophore concentration, this result establishes time-resolved measurements as more reliable than intensity-based approaches for collagen characterization. Practically, our data identify 355 nm excitation as offering the best balance between excitation efficiency and photobleaching resistance for future fluorescence-based collagen studies.

### 4.4 Study Limitations and Future Directions

This study establishes reference fluorescence signatures for purified collagen I–V, but a few limitations are noted. First, the I-EEM and TR-EEM acquisition involves extended illumination, which can induce photobleaching (especially early in the scan) and bias intensity-based spectra. Future work should reduce light dose and temporal bias by shortening dwell times and optimizing or randomizing wavelength acquisition order, and by adopting more photon-efficient acquisition strategies.

Second, measurements on purified collagen samples under simplified hydration conditions cannot fully represent native tissue complexity, including fibril organization, heterogeneous crosslinking, and mixed fluorophore backgrounds. Future studies should validate these findings in ECM phantoms and ex vivo/in vivo tissues under more physiological conditions and quantify how tissue structure and optics affect both spectral and time-resolved metrics.

Finally, time-resolved decays were reconstructed using Laguerre expansion–based deconvolution, which provides a model-agnostic estimate of the fluorescence decay. While we emphasized average lifetime for robust comparison and consistency with typical clinical FLIm workflows, the reconstructed decays contain additional information that can be exploited in future analyses, such as Laguerre coefficients, phasor features, and multi-exponential approximations.

## 5 Conclusion

This study establishes the first comprehensive I-EEM and TR-EEM reference dataset for collagen Types I–V. Three principal findings emerge: (1) both collagen type and organ source independently shape fluorescence signatures, necessitating organ-specific reference for accurate fluorescence signature characterization; (2) hydration consistently red-shifts emission peaks and reduces fluorescence lifetimes, highlighting that in vivo interpretation should rely on hydrated reference measurements rather than dry-only benchmarks, and (3) fluorescence lifetimes remain stable under photobleaching, establishing time-resolved approaches as the more robust modality for quantitative collagen assessment. This dataset and the associated findings provide a foundation for optimizing fluorescence-based clinical diagnostics, tissue engineering applications, and future translational research.

## Code and Data Availability

The data that support the findings of this study are available from the corresponding author upon reasonable request.

## Acknowledgements

This work was supported by the National Institutes of Health (Grants No. P41EB032840, No. R01CA187427, No. R01HL157712, No. R01CA250512). The authors acknowledge the use of a large language model (Claude) for language editing and grammar refinement of this manuscript.

